# SeekNSee: A Rule-Based Framework for Prioritizing Proteotypic Peptides

**DOI:** 10.64898/2026.11.07.764955

**Authors:** Mauricio Frota Camacho, Guilherme Pauperio Lanfredi, Adriana Franco Paes Leme, Nilson Antônio da Rocha Coimbra, Jackson Gabriel Miyamoto

**Affiliations:** Advanced Clusters in Health Technologies (NATS), Brazilian Biosciences National Laboratory (LNBio), Brazilian Center for Research in Energy and Materials (CNPEM), Campinas, São Paulo, Brazil; Graduate Program in Pharmaceutical Sciences (PPG – FCF), Universidade Estadual de Campinas (UNICAMP), Campinas, São Paulo, Brazil; Departament of Basic and Oral Biology, Faculty of Dentistry of Ribeirão Preto, University of São Paulo (FORP/USP), Ribeirão Preto, São Paulo, Brazil

**Keywords:** targeted proteomics, peptide prioritization, proteotypic peptides, in silico digestion, rule-based scoring, traceability, R/Shiny, DIA, DDA, bioinformatics software

## Abstract

Selecting peptides candidates for target proteomics requires integrating a multicriteria problem, particularly when users lack prior experimental data, programming expertise, or stable access to external repositories. We developed SeekNSee, a locally executed R/Shiny application designed to improve the reproducibility and auditability of peptide-candidate screening through enzyme-aware in silico digestion, configurable and traceable rule-level decision tracking. Starting from a user provided reference proteome and target sequences, SeekNSee generates peptides-candidates using nine digestion configurations and evaluates them with 12 rules configured as *ignore*, *apply*, or *block*. Numerical score and retained/blocking status are stored independently, and every candidate-rule outcome is preserved in a rule-level trace. Computational validation confirmed the specified behavior in controlled cases through 198 passing automated assertions, complemented by successful operational and processing tests on Windows and Ubuntu. Proteome-wide application covered all 20,416 reviewed human proteins, generating 733,864 peptide-candidate and 581,026 unique peptides sequences while preserving data integrity, decision traceability, and candidate-specific context. DIA, DDA, and deep-fractionated proteomics benchmarking showed strong associations between SeekNSee scores categories, peptide observation, and multiple analytical measurements. Together, these results support that SeekNSee provides a transparent and traceable screening layer for generating peptide-candidates for subsequent assay and experimental validation.

## 1. INTRODUCTION

Targeted mass spectrometry methods, including SRM/MRM and PRM, enable sensitive and reproducible measurement of predefined proteins through selected surrogate peptides. Their analytical performance therefore depends on choosing candidates that represent the proteins of interest and are compatible with sample preparation, chromatography, and mass-spectrometric acquisition ^1,2^. Accordingly, peptide selection constitutes a critical and multicriteria step in the development of targeted proteomics assays rather than a secondary consequence of instrument configuration ^1–3^. Candidate prioritization may be influenced by sequence specificities within the relevant proteome and characteristics that can represent different biological and analytical dimensions and do not provide equivalent evidence of peptide suitability ^2,3^.

Computational strategies integrating empirical evidence and rule-based criteria have emerged to navigate the complex landscape of proteotypic peptide selection, supporting systematic, transparent, and reproducible prioritization ^2–4^. Rule-based tools have sought to standardize this process. PeptidePicker ^3^ combines information from multiple databases with weighted filters, Typic ^2^ integrates established criteria with public repositories and user-provided DDA results, and PepMANDIS ^4^ evaluates public or user-defined protein collections using sequence-derived properties.

Predictive approaches have additionally modeled peptide detectability from sequence-derived and experimentally learned features ^5^. PeptideRanger uses a random forest classifier and supports experiment-specific retraining ^6^, whereas AP3 incorporates predicted digestibility ^7^, and DeepDetect extends detectability modeling across multiple proteases ^8^. These studies emphasize that experimental observation depends both on peptide generation and on subsequent analytical behavior ^9–11^. Protease identity further changes cleavage boundaries, peptide properties, and sequence coverage, and even the conventional exclusion of tryptic K/R–P cleavage is not absolute ^12^. Although increasingly sophisticated, predictive scores remain dependent on their training data, feature representation, and experimental context ^5–8^.

Experimentally derived resources provide complementary evidence for candidate selection. DDA results, targeted-assay repositories, and spectral libraries document previous observations and support transition selection, retention-time evaluation, and assay development ^2,13,14^. Their reuse nevertheless depends on resource coverage, metadata quality, and the biological and analytical context of the original measurements, limiting their applicability to novel sequences, poorly characterized organisms, alternative proteases, or projects without prior discovery data ^2,4,13,14^.

Existing tools therefore address overlapping but distinct aspects of candidate selection and differ in their dependence on experimental evidence, predictive models, spectral resources, external repositories, and user-defined databases ^2–8,13,14^. These requirements are appropriate for many applications but can create an operational gap during early assay planning, when users may possess only target sequences and a reference proteome and may lack prior experimental data or the expertise required to construct a custom computational workflow ^2–4,6–8,13^.

Reducing this operational barrier is insufficient if an analysis cannot be reproduced and its decisions reconstructed. Computational reproducibility requires preservation of software versions, input data, dependencies, parameters, intermediate operations, and workflow provenance ^15,16^. For peptide prioritization, reproducibility additionally requires recording how candidate-specific evidence was converted into ranking and classification decisions ^15–17^. An accessible exploratory framework should therefore support both execution and reconstruction of the analytical context and decision process. This requirement exposes an interpretative gap. Predictive scores estimate detectability but do not necessarily provide a customized profile of which specificity-, sequence-, position-, or digestion-related criteria support or oppose an individual candidate ^5–8^. Likewise, a fixed ranking may not represent the policies required for different analytical objectives ^2,3,6^. Users may wish to treat the same criterion as a preference, an exclusion condition, or contextual information. A transparent decision-support framework should therefore preserve both the result of each criterion and the policy used to interpret it ^18^.

To address these operational and interpretative gaps, we developed SeekNSee, a local R/Shiny application for exploration in silico digestion and traceable rule-based prioritization of peptide-candidates. By encoding empirical knowledge in explicit peptide-selection rules, SeekNSee considers sequence- and context-derived characteristics relevant to peptide observation, identification, and quantification without requiring prior experimental spectra or model training on labeled peptide-detectability datasets. The framework integrates enzyme-aware candidate generation, user-defined reference-proteome specificity, configurable rules states, and numerical scores within a locally executed-graphical workflow. Users can configure and inspect prioritization decisions without writing code, modifying command-line scripts, or querying repositories in real time.

Here, we present SeekNSee and evaluate the scientific and operational behavior of its candidate-prioritization framework. Versioned computational tests assess consistency with the specified digestion and decision logic, while proteome-wide processing of the reviewed human proteome evaluates peptide-candidate generation, coverage, and traceability. Complementary DIA and DDA data sets are used to examine associations between SeekNSee scores, peptide observation, identification confidence, and quantitative measurements. Finally, PeptideRanger provides a comparison with a machine-learning-based detectability predictor, and an independent deep-fractionated data set evaluates whether score-observation associations persist under increased proteome coverage.

## 2. EXPERIMENTAL SECTION

### 2.1. Implementation and Availability

SeekNSee was implemented in R using Shiny and distributed as a self-contained application directory containing the components required for local execution. The graphical interface is opened in the default web browser and allows users to configure analyses, inspect peptide-candidates, modify rule policies, and export complete results without writing code or editing command-line scripts. The software is available at https://github.com/camachomf/SeekNSee.

### 2.2. Software Architecture and Processing Workflow

SeekNSee organizes the analysis into a processing workflow that links sequence input, peptide-candidate generation, rule evaluation, and result inspection and export. FASTA records are parsed into normalized sequence objects while retaining original identifiers and annotations. The selected enzyme model defines cleavage boundaries, from which candidate instances across are generated in the permitted missed-cleavage range. Each candidate is rated against all 12 rules, with applicability, pass/fail outcome, score contribution, blocking outcome, and rule-specific evidence recorded for each evaluation. Candidate-level results are subsequently organized for graphical inspection and structured Excel export.

The same scientific processing implementation is used across platform-specific distributions, while launchers, runtime packaging, and operating-system integration are maintained separately. The complete processing workflow, representative interface components, and candidate-level traceability are illustrated in Figure 1.

**Figure 1.**
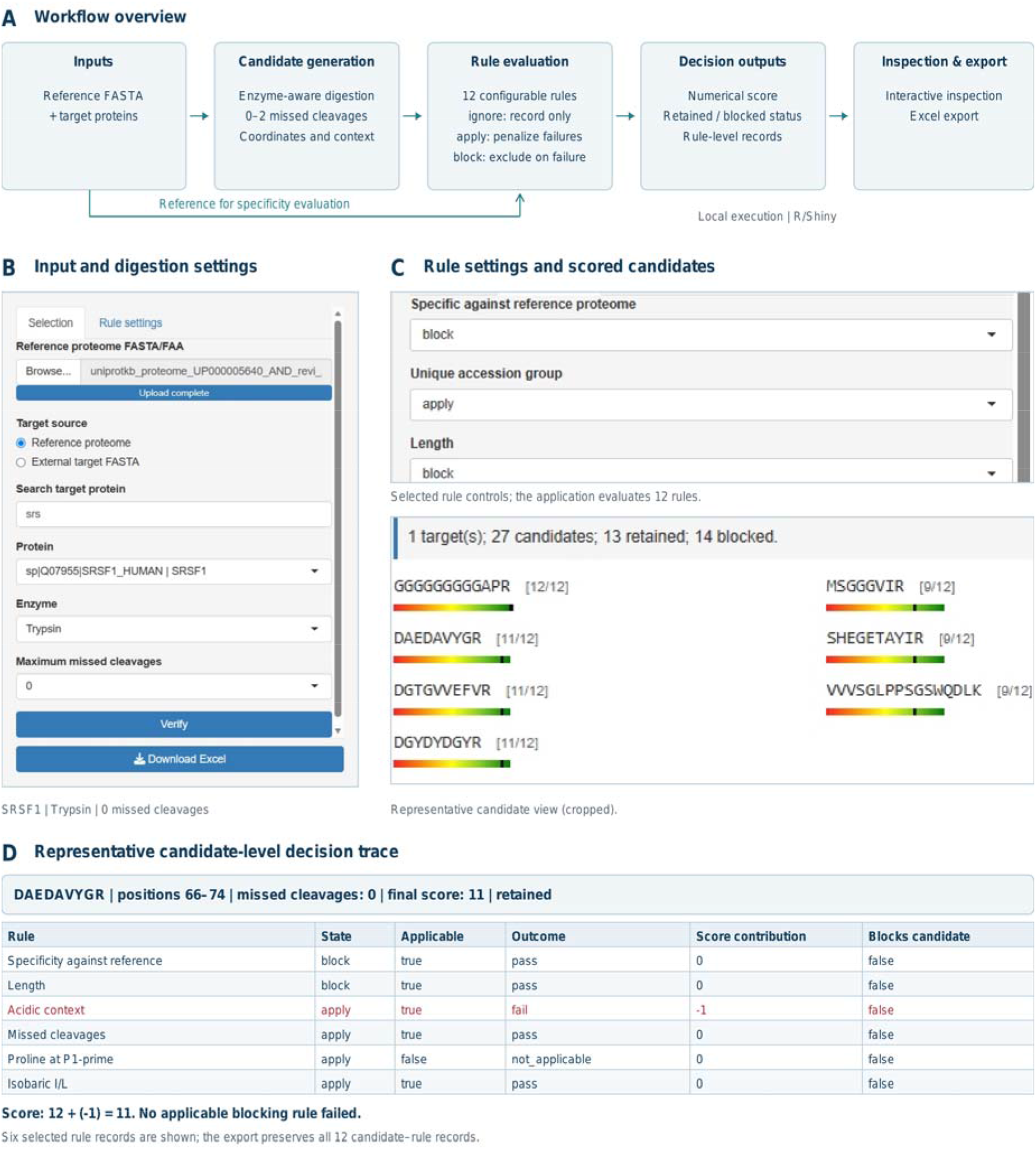
SeekNSee workflow, graphical interface, and rule-level traceability. (A) Sequence inputs, enzyme-aware candidate generation, configurable rule evaluation, and inspection and export. The reference FASTA defines the context for specificity assessment. Numerical scores and retained/blocked status are recorded separately. (B) Cropped screenshot showing SRSF1 selection and digestion settings. (C) Cropped screenshot showing selected rule controls and scored candidates; only part of the interface is displayed. (D) Schematic table based on exported records of six exported rule records for “DAEDAVYGR” (positions 66-74). The acidic-context failure contributes “-1”, yielding a score of 11; no applicable blocking rule fails. The complete export preserves all 12 candidate-rule records.

### 2.3. Input Sequences and Candidate Generation

SeekNSee requires a reference proteome in FASTA format and one or more target-protein sequences selected from the reference file or supplied externally. External targets are processed according to the selected reference-specificity policy. The principal inputs, core operations, and outputs of the SeekNSee workflow are summarized in Table 1.

**Table 1.** Inputs, core operations, and outputs of the SeekNSee workflow.

| **Input** | **Core operation** | **Output** |
|---|---|---|
| Reference proteome in FASTA/FAA format | Parse and standardize sequences, retain identifiers, and define the reference universe for specificity assessment. | Reference sequences and identifiers defining the analysis-specific sequence context. |
| Target proteins selected from the reference proteome or supplied in a separate FASTA/FAA file | Read target sequences and retain their identifiers and available annotations. | Target-protein records linking generated candidates to their source sequences. |
| Enzyme selection and maximum missed-cleavage allowance (0–2) | Apply enzyme-specific cleavage definitions and generate candidate instances within the configured missed-cleavage limit. | Candidate sequences with target identifiers, coordinates, missed-cleavage counts, and cleavage context. |
| State assigned to each of 12 rules: ignore, apply, or block | Evaluate applicability and pass/fail outcomes; calculate score penalties and apply exclusion criteria independently. | Numerical score, separate retained/blocked status, and rule-level records of settings, outcomes, penalties, and blocking reasons. |
| Completed analysis and associated run configuration | Organize candidate-level and rule-level results for interactive inspection and structured Excel export. | Ranked and blocked candidate tables, target records, run metadata, rule configuration, and complete candidate-rule trace records. |

The application supports nine digestion configurations: Trypsin, Trypsin/P, LysC, ArgC, AspN, GluC, Chymotrypsin, Elastase, and ProAlanase. Enzyme-specific cleavage boundaries are used to generate candidates containing zero, one, or two missed-cleavage events, according to the user’s selection. Candidates of 4–25 amino acids are retained for evaluation, whereas the Length rule independently assesses the preferred interval of 8–25 residues. Each candidate remains linked to its target protein, sequence coordinates, and cleavage context. Complete cleavage definitions and enzyme-dependent rule applicability are provided in Table S1.

### 2.4. Rule-Based Prioritization and Traceability

Each candidate starts with a score of 12 and is evaluated against 12 sequence- and context-based rules. Complete rule definitions, applicability conditions, scoring effects, blocking behavior, trace and linked evidence fields, and the configuration used in the empirical acceptance analysis are provided in Table S2. Users configure each rule as *ignore*, *apply*, or *block*. Ignore rules are evaluated and recorded without affecting the score or candidate status. Under apply, each failed rule subtracts one point, except “Missed cleavages”, which subtracts one point per missed cleavage. Under block, a failed rule excludes the candidate from the retained set without directly changing its numerical score. Evaluation continues after a blocking event so that all exclusion reasons are retained. Nonapplicable rules neither change the score nor block candidates. Their scores and status are stored separately, and rule-level trace records each criterion, its configuration, and contribution to the decision.

Accordingly, the numerical score for candidate is calculated as

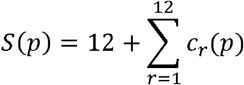

where *c_r_(p)* is the contribution of rule *r*. Ignored, nonapplicable, passed, and blocking rules contribute 0 to the numerical score. Standard failed rules in apply contribute −1, and the applied Missed cleavages rule contributes the negative number of missed-cleavage events.

Analysis results are exported as a structured Excel workbook containing the Run_summary, Rule_configuration, Target_proteins, Ranked_peptides, Rule_trace, Blocked_peptides, and Rule_definitions worksheets. Stable target_id, candidate_key, and rule identifiers link target-, candidate-, and rule-level information across the workbook. Worksheet purposes, literal column names and order, data types, permitted values, examples, relational keys, missing-value conventions, and long-cell abbreviation rules are described in Table S3.

### 2.5. Computational Validation and Proteome-Wide Processing

Computational validation assessed digestion, candidate generation, rule evaluation, scoring, blocking, rule-trace construction, export behavior, and internal consistency against predefined expectations. Proteome-wide scalability was evaluated using the frozen SeekNSee 0.2.5 scientific core and default 12-rule configuration with Trypsin, zero missed cleavages, and candidate lengths of 4–25 amino acids across the 20,416-entry reviewed human reference proteome. Processing was conducted in 205 deterministic batches, followed by independent auditing of scientific identity, proteome coverage, count consistency, transactional integrity, and file identity. Candidate instances were then aggregated and deduplicated by exact peptide sequence while preserving occurrence-level provenance.

Computational validation and packaging were conducted for Windows 10/11 x64 and Ubuntu LTSx86_64 distributions, and platform-specific tests are summarized in Supplementary Table S4, whereas the complete proteome-wide audit and sequence-level analyses are reported in Supplementary Table S5.

### 2.6. Experimental Data Sets and LC-MS/MS Acquisition

Experimental evaluation used DIA and DDA analyses of the Pierce HeLa Protein Digest Standard (Thermo Fisher Scientific). DIA measurements used 200 ng of digest in five technical replicates, whereas DDA used 100 ng in three technical replicates. The two data sets provided complementary acquisition conditions for evaluating candidate prioritization.

DIA data were acquired on Orbitrap Astral coupled to Vanquish Neo UHPLC and FAIMS Pro Duo interface. Detailed chromatography, acquisition parameters, and database search results were deposited in the ProteomeXchange Consortium via the PRIDE repository (PXD085697).

### 2.7. Proteomics Data Processing

DIA data were processed with DIA-NN 2.5.1 ^19^ using a predicted spectral library generated from the reviewed canonical human reference proteome (UniProt UP000005640) ^20^, containing 20,416 entries. DDA data was processed with MaxQuant against the same reference sequences. Both workflows used trypsin specificity, up to two missed cleavages, fixed carbamidomethylation of cysteine, no variable modification, and 1% false discovery rate threshold at precursor level.

Peptide-level features were extracted from the DIA-NN (*report.parquet*) and MaxQuant (*evidence.txt*) outputs after contaminant filtering. Data filtering, aggregation, and visualization were performed in R (4.4). Only precursor features detected in all technical replicates of the corresponding experiment were retained. Precursor features were averaged across replicates. Charge-states were collapsed by stripped peptide sequence, and the precursor with the highest mean abundance was retained as the representative feature for each stripped sequence.

Experimental peptide sequences were matched exactly to the theoretical candidates. Peptides meeting the replicate-consistency criterion were classified as observed for the evaluation; unmatched candidates were classified as not meeting the criteria, rather than as intrinsically undetectable.

### 2.8. Experimental Evaluation and PeptideRanger Comparison

Experimental evaluation used a fixed SeekNSee rule configuration: “Reference-proteome specificity” and “Length” were configured as block, “Accession-group uniqueness” and “Isobaric I/L” as ignore, and the remaining applicable rules according to the configuration reported in the Table S2. The same configuration was used across all experimental evaluations.

SeekNSee and PeptideRanger version 0.1.0 were evaluated using theoretical candidate set, with matched digestion and sequence-processing settings. Peptides satisfying the experimental observation criterion constituted the observed class, whereas the remaining candidates in the shared theoretical universe constituted the comparison class. Receiver-operating-characteristic (ROC) analysis assessed the extent to which each score ranked experimentally observed peptides above the remaining candidates.

Spearman rank correlations were used to examine associations between the prioritization scores and peptide-level identification confidence, chromatographic properties, and quantitative measurements. Physicochemical properties were calculated using the Peptides R package ^22^. Evaluated variables, source columns, transformations, missing-value treatment, sample sizes, statistical procedures, package versions, and software-environment information are provided in Supplementary Methods and Table S6.

### 2.9. Evaluation Using an Independent Deep-Proteome Data Set

An independent evaluation used the public HEK293 data set PXD046372 23 to examine whether associations between prioritization and peptide observation persisted under increased proteome coverage. The data set comprised 46 offline high-pH reversed-phase fractions analyzed in triplicate by narrow-window DIA. Data were processed with DIA-NN, and peptide sequences were mapped to the same theoretical candidate representation used in primary evaluation. To produce a non-redundant peptide-level dataset, the same collapsing strategy was applied, retaining the fraction with the highest precursor abundance as the representative observation.

## 3. RESULTS AND DISCUSSION

### 3.1. Computational Validation and Traceable Proteome-Wide Application of SeekNSee

Computational and operational testing of SeekNSee 0.2.5 confirmed that the implementation behaved as specified across the evaluated configurations, with all 198 automated assertions passing, while manual and package-level confirmed consistent behavior of the validated scientific core across Windows 10/11 x64 and Ubuntu LTSx86_64 distributions. Proteome-wide processing covered all 20,416 reviewed human reference proteome entries in 205 independently audited batches, with no missing or duplicated protein indices or identifier/sequence mismatches. It generated 733,864 candidate instances (384,954 retained; 348,910 blocked) and 8,806,368 rule-trace records; 19,859 proteins had at least one retained candidate, whereas 557 had none under the evaluated default configuration. Exact-sequence deduplication yielded 581,026 unique peptides (383,717 consistently retained; 197,309 consistently blocked), with no status conflicts but instance-level score variability for 10,629 sequences.

Together, these results support the operational consistency of SeekNSee and the preservation of decision traceability during proteome-wide processing under the evaluated configuration. Although retained/blocked status was consistent across instances of each peptide sequence, instance-level score variability highlights the importance of preserving protein associations and candidate context rather than interpreting sequence-level scores in isolation. Supplementary Table S4 summarizes the validation framework; Supplementary Table S5 reports batch integrity, protein coverage, aggregate distributions, sequence deduplication, status conflicts, recovery provenance, and external-artifact identities.

### 3.2. SeekNSee Scores Association with Analytical Features in DIA Data

We performed an empirical evaluation of peptide performance using confidence and observability metrics to encompass identification reliability and quantitative robustness within next generation mass spectrometry proteomics workflows ^18^. HeLa cell digest was selected as the primary benchmark proteome due to its extensive characterization and routine use as a standard sample in proteomics studies, providing a well-established framework for evaluating peptide observability and analytical performance ^10^.

The first exam was using a HeLa digest analyzed by DIA on an Orbitrap Astral. After DIA-NN processing and replicate-level filtering, 117,094 precursors were identified, of which 81,674 were mapped to the theoretical candidate universe. The corresponding theoretical space contained 1,671,645 sequences derived from 20,416 entries of the canonical human proteome. Experimentally observed sequences were distributed across eight SeekNSee score categories, with higher categories displaying a greater proportion of observed peptides than the complete theoretical sequence space (Figure 2A). This pattern suggests that the rule-based score contains information related to contextual peptide observation, despite not having been trained as a detectability classifier. Nevertheless, the comparison does not establish that unmatched theoretical sequences are intrinsically undetectable and should be interpreted as discrimination in this specific dataset. Lack of observation may reflect protein abundance, sample composition, instrument acquisition, database-search, modification state, or other forms of experimental censoring ^18^.

**Figure 2.**
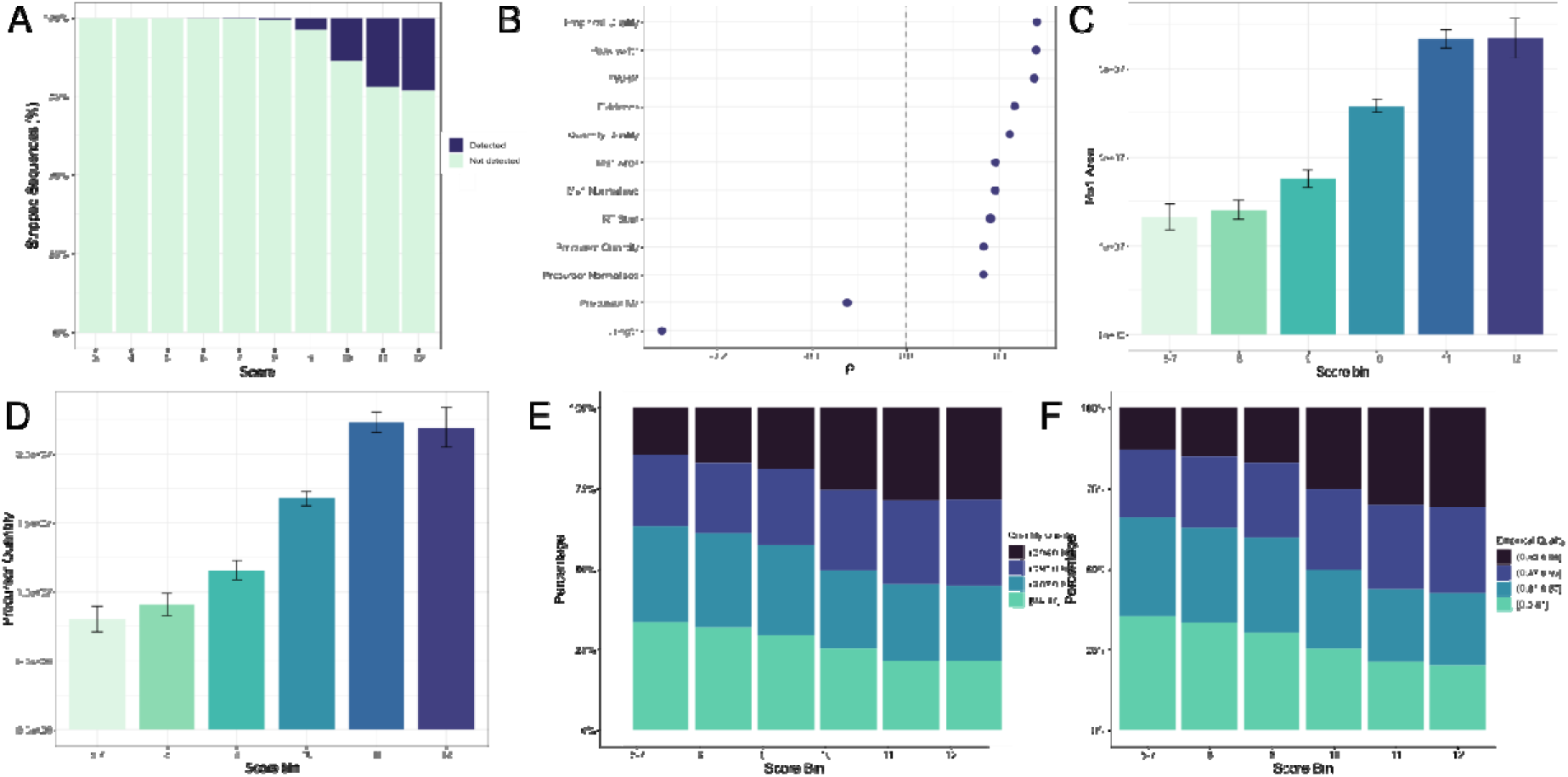
Evaluation of SeekNsee scoring for peptide quality and detection. HeLa DIA acquisitions were used to assess the performance of the SeekNsee scoring framework and its relationship with peptide analytical metrics. (A) Distribution of identified proteotypic peptide sequences across score categories. (B) Spearman correlation coefficients (ρ) between scores and DIA-NN-derived features. Average MS1 (C) and MS2 (D) precursor abundance across score categories, with standard errors of the mean. Percentage distribution of Quantity Quality (E) and Empirical Quality (F) binned score across SeekNsee score categories.

Correlation analysis revealed that the SeekNsee score was positively associated with peptide abundance, increased chromatographic peak dimensions, later chromatographic elution, and improved quantitative quality metrics, whereas precursor *m/z* and length displayed negative correlations (Table S6 and Figure 2B). Higher-scoring candidates exhibited broader chromatographic peak dimensions (Figure S1) and greater MS1- and MS2-level abundance (Figure 2C-D), which may contribute to more consistent peak detection than low-abundant signals. Systematic differences in DIA-NN derived scores categories indicate lower estimated precursor instrumental quantification error and coherent agreement between quantitative features of precursors and transitions (Figures 2E-F and illustrated by Figure S2), associated with scoring.

DIA-NN ranks calculates a weighted discriminant score based on the concordance of multiple orthogonal features that provide a quantitative measure of signal quality and reliability, representing a summarization of several underlying correlation measurements ^19^. By adopting DIA as the primary evaluation strategy, we captured a broader dynamic range of peptide observations while reducing the dependence of detectability assessment on binary classification based solely on the presence or absence of precursor-level signal ^18^. Evaluating the distribution of these features across identified peptides enables the assessment of identification robustness and supports the selection of high-confidence identifications ^21^. These metrics are particularly relevant when selecting candidates for targeted experiment based on empirical discovery analysis, allowing for flexibility of quantitation levels, stable covariation of measured signals and ensuring direct correlation of analyte abundance and monitored identity fragments. Thus, we leveraged the software’s integrated scoring metrics to assess peptide proteotypicity and analytical suitability.

### 3.3. Complementary DDA Analysis

Consistent with DIA analysis, the complementary DDA dataset exhibited similar trends across the evaluated parameters (Figure 3A). Precursor intensity (Figure 3B) and Andromeda score (Figure 3C) were positively associated with peptide classification. As a probabilistic measure of peptide-spectrum match quality, the Andromeda score increased across SeekNsee score categories, indicating enrichment of highly ranked peptides with confidently identified peptide-spectrum matches ^22^. Identified stripped peptide sequences exhibited stronger negative correlations with molecular mass (ρ = −0.34; Figure 3D), peptide length (ρ = −0.26), and retention time-related features, including chromatographic elution and MS detection (ρ = −0.13; Figure 3E and Table S6), than observed in the DIA dataset.

**Figure 3.**
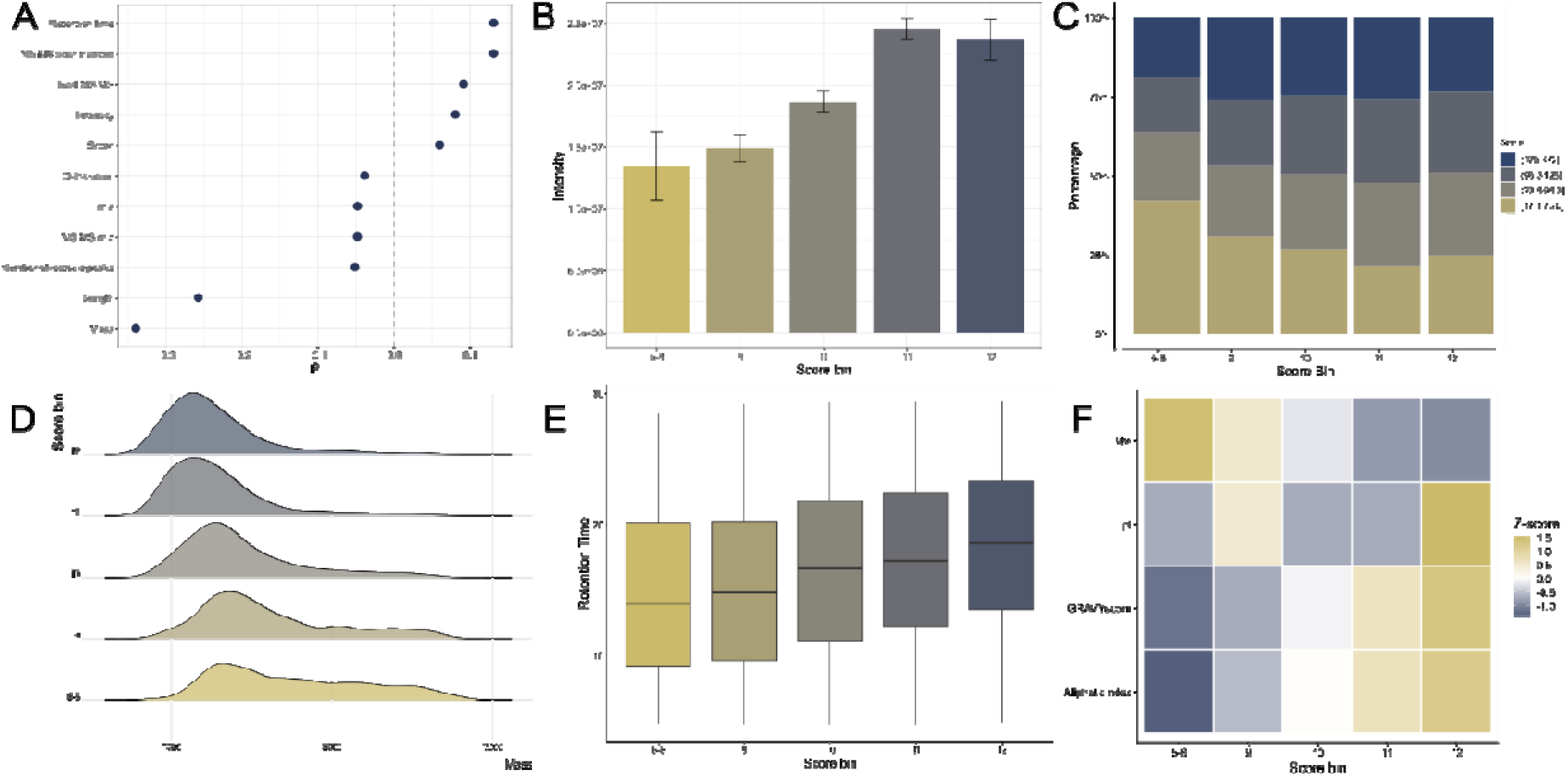
Evaluation of SeekNsee scoring in a complementary DDA dataset. (A) Spearman correlation coefficients (ρ) between SeekNsee scores and MaxQuant-derived features. (B) Mean precursor intensity across score categories with standard errors of the mean. (C) Percentage distribution of binned Andromeda scores across SeekNsee score category categories. (D) Peptide mass density distribution across score categories. (E) Retention time distributions represented as boxplots across score categories. (F) Heatmap showing Z-scores of peptide physicochemical properties.

To further investigate these associations, we evaluated the physicochemical properties of the identified sequences (Table S6). Higher SeekNsee score categories were enriched for peptides with increased isoelectric point (pI), hydrophobicity (GRAVY), and aliphatic index (Figure 3F). Top-scoring sequences tend to be shorter, more hydrophobic, and more basic. These patterns are plausible consequences of the rule architecture that contribute directly or indirectly to prioritization, preferentially ranking peptides that exhibit physicochemical characteristics associated with favorable chromatographic retention and peptide observability in LC-MS workflows ^23,24^.

### 3.4. Comparision of SeekNsee and PeptideRanger

We next benchmarked SeekNsee against PeptideRanger, a random forest-based predictor of peptide-detectability trained on the human proteome using UniProt/Swiss-Prot canonical sequences ^6^. The tool was selected owing to its comprehensive evaluation against other peptide detectability prediction tools, providing a well-characterized reference framework for comparison. In the current study, PeptideRanger and SeekNSee achieved ROC-AUC values of 0.89 and 0.84, respectively, in the HeLa DIA dataset (Figure 4A and Table S7). Corresponding values of 0.90 and 0.88 were reported for the complementary DDA dataset (Figure 4B). This difference is expected, as PeptideRanger relies on a continuous random forest model, whereas SeekNSee employs a simple discrete scoring framework. Nevertheless, the observed performance is comparable to that reported for contemporary machine learning and deep learning-based predictors as reviewed by Schilling ^18^. Furthermore, both score systems showed associations with several peptide-level analytical measurements, although the direction and magnitude of these associations differed between frameworks (Figure 4C-D, Tables S6 and S8).

**Figure 4.**
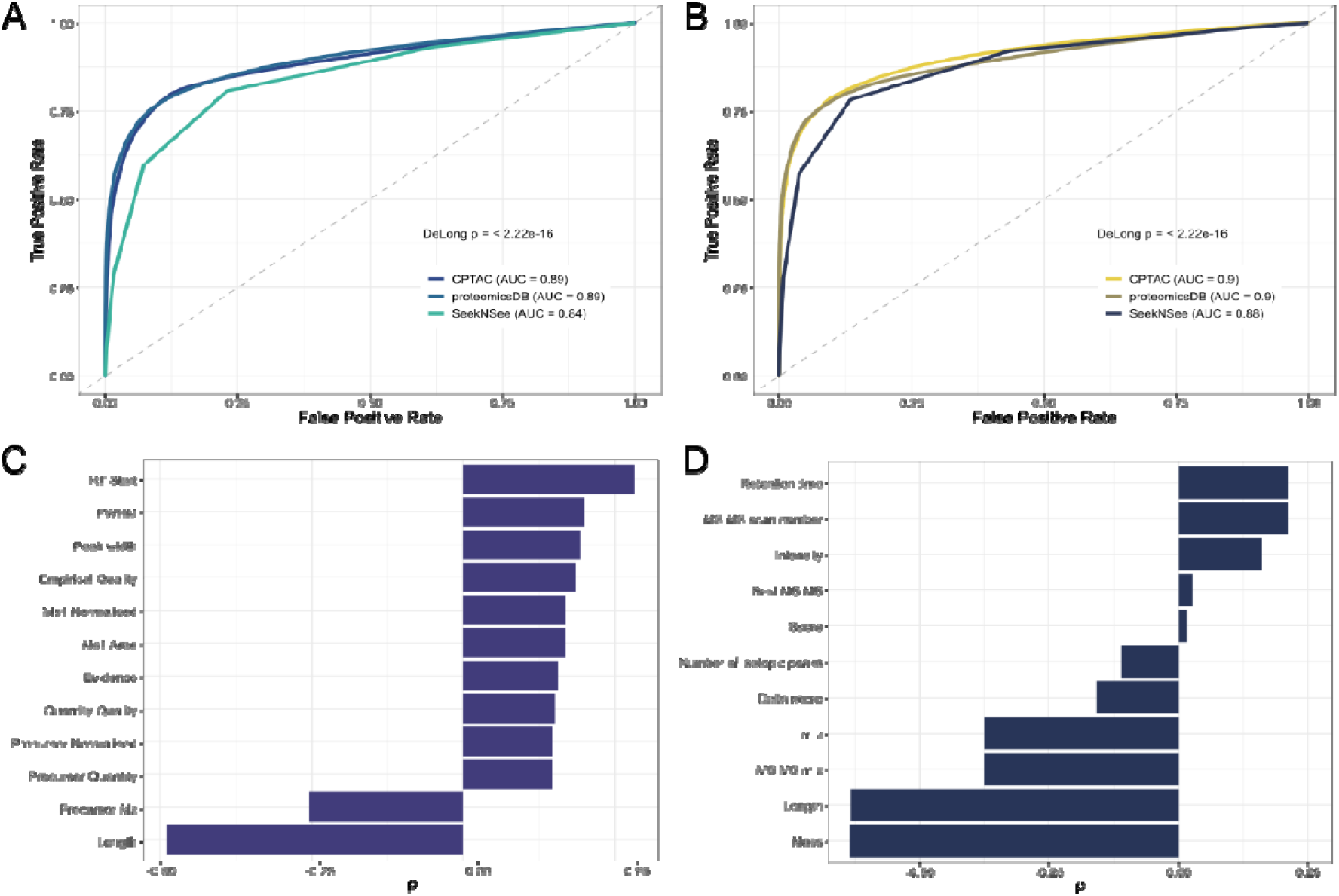
Benchmarking of SeekNsee against the machine learning-based peptide detectability predictor PeptideRanger. (A) Receiver operating characteristic (ROC) curves comparing SeekNsee and PeptideRanger in the HeLa DIA dataset. (B) ROC curves comparing SeekNsee and PeptideRanger in the HeLa DDA dataset. Correlation of peptide identification confidence metrics across PeptideRanger random forest score categories in the DIA (C) and DDA (D) datasets.

### 3.5. Deep Proteome Coverage Expanded Peptide Detection

An independent deep-proteome analysis using the public HEK293 dataset ^25^ (PXD046372) was performed to examine whether the relationship between score and observation in a larger fraction of the theoretical peptide space sampled experimentally. Comprising 46 offline high-pH reversed-phase fractions, the reanalysis reported 357,353 precursor identifications and 12,450 protein groups after DIA-NN processing.

The increased proteome coverage substantially improved precursor detection, with the highest SeekNsee score categories encompassing nearly half of all theoretical peptide sequences generated from the canonical human proteome (Figure 5A). A comparable distribution was observed for PeptideRanger classifications (Figure 5B). Despite similar detectability profiles, SeekNsee scores presented a weak yet significant monotonic relations with lower q-values and PEP (Figure 5C, E), whereas higher PeptideRanger scores showed the opposite trend, accompanied by increased interquartile dispersion of both metrics (Figure 5D, F). These findings indicate that peptides prioritized by SeekNSee exhibit subtle consistency in identification confidence and lower estimated local and global error rates q-values and PEP provide complementary measures of global and local precursor level identification confidence 26. The systematic divergence between the scoring frameworks suggests that the SeekNsee heuristic captures additional aspects of analytical performance beyond peptide detectability alone.

**Figure 5.**
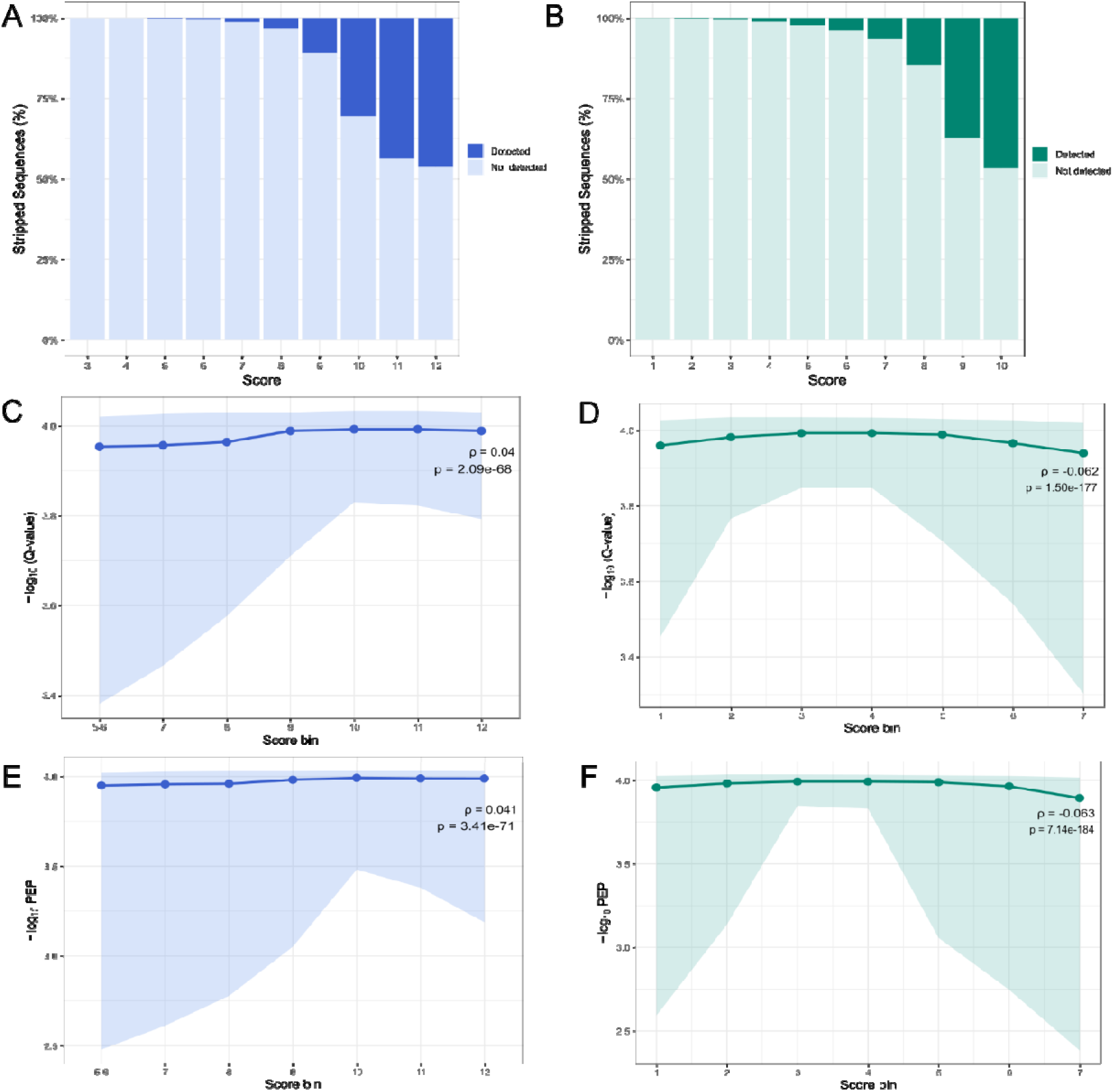
Comparison of SeekNsee and PeptideRanger in an independent deep-proteome dataset. The HEK293 dataset was used to assess peptide detectability and identification confidence under increased proteome coverage. (A-B) Percentage of theoretical peptide sequences detected across SeekNsee and PeptideRanger score categories. (C-F) Median and interquartile range of −log10(PEP) across SeekNsee and PeptideRanger score categories, respectively. Spearman correlation coefficients (ρ) and corresponding p-values are shown in each panel. To facilitate direct visual comparison between frameworks, PeptideRanger random forest scores were binned into quantiles corresponding to the SeekNsee score categories.

By integrating empirical knowledge embedded within established peptide-selection rules, the framework prioritizes peptide-candidates that are not only observable but also reliably identified and quantified across different proteomic datasets, acquisition strategies, instruments, and data-processing workflows. These findings highlight the ability of SeekNsee to rank peptide-candidates in silico without requiring experimental spectra or large model training on labeled peptide detectability datasets.

### 3.6. Integrated Interpretation and Limitations

Across DIA, DDA, and independent deep-proteome evaluations, higher SeekNSee scores were associated with peptide observation, identification confidence, quantitative signal, and selected chromatographic and physicochemical features, supporting the framework’s empirical relevance without establishing a calibrated detection probability or guaranteeing targeted-assay performance. Associations with peptide length, basic-residue content, and sequence motifs may partly reflect their direct or indirect contributions to scoring. Observation also depends on source-protein abundance, sample composition, digestion efficiency, ionization, chromatography, acquisition, database-search settings, modifications, detection limits, and stochastic or acquisition-dependent sampling; unmatched theoretical candidates are therefore not intrinsically undetectable. Comparisons with PeptideRanger do not establish superiority because learned detectability prediction and configurable, traceable rule-based prioritization represent distinct, complementary paradigms requiring context-specific experimental validation.

## 4. CONCLUSION

SeekNSee provides an accessible, configurable, and traceable framework for peptide-candidate prioritization in targeted proteomics. By combining user-defined reference proteomes, enzyme-aware in silico digestion, and explicit sequence- and context-based criteria within a locally executed graphical interface, the software reduces the theoretical candidate space without requiring prior experimental spectra or real-time access to external repositories. Versioned computational tests supported the deterministic behavior and internal consistency of the evaluated scientific core, and proteome-wide processing demonstrated complete coverage of the reviewed human reference proteome with auditable candidate- and rule-level provenance. DIA, DDA, and deep-proteome benchmarking further showed associations between SeekNSee score categories, peptide observation, and multiple identification, quantitative, chromatographic, and physicochemical measurements. SeekNSee is not intended as a calibrated predictor of peptide detectability or a substitute for empirical assay validation. Instead, it provides a reproducible decision-support layer for comparing selection policies, documenting why candidates were retained, penalized, or blocked, and generating experimentally actionable peptide lists for subsequent validation.

## ASSOCIATED CONTENT

**Supporting Information** (.xlsx)

Table S1. Detailed enzyme definitions.

Table S2. SeekNsee rule definitions and applicability.

Table S3. Software validation summary.

Table S4. Computational provenance and reproducibility information.

Table S5. Additional software implementation details.

Table S6. Correlation analysis of experimental datasets.

Table S7. ROC-AUC analysis of peptide detectability.

Table S8. Correlation analysis of PeptideRanger random forest scores.

**Supplementary Information** (.docx)

Figure S1. Relationship between chromatographic peak dimensions and precursor quantification metrics.

Figure S2. Extracted ion chromatogram of ISSPTETER.

## AUTHOR INFORMATION

Requests for further information should be directed to the corresponding author, Maurício Frota Camacho.

## AUTHOR CONTRIBUTIONS

Maurício Camacho: original draft, Visualization, Methodology, Investigation, Writing – original draft, Formal Analysis and Data Curation. Guilherme Lanfredi: Methodology, score strategy and graphical user. Nilson Coimbra: software e data management. Adriana Paes Leme: Grant management, Writing – original draft. Jackson Miyamoto: benchmarking, data analysis, original draft, Visualization, Methodology, Investigation, Writing – original draft, Formal Analysis and Data Curation. All authors reviewed, revised, and approved the final manuscript.

## ACKNOWLEDGMENTS

This work was supported by Conselho Nacional de Desenvolvimento Científico e Tecnológico grants 380453/2026-6 to J.G.M. and 80454/2026-2 to G.P.L., Fundação de Amparo à Pesquisa do Estado de São Paulo 2021/06125-6 to MFC, EMU 2022/11476-5 and 2018/18496−6 to A.F.P.L and CNPEM-USP 2026-824 to NARC. This study was conducted at the Mass Spectrometry Laboratory of the Brazilian Biosciences National Laboratory (LNBio) and supported by and had resources from the Brazilian Federal Government provided to the Brazilian Center for Research in Energy and Materials (CNPEM), a private nonprofit organization under the supervision of the Brazilian Ministry for Science, Technology, and Innovation (MCTI).

## DECLARATION OF GENERATIVE AI AND AI-ASSISTED TECHNOLOGIES IN THE MANUSCRIPT PREPARATION PROCESS

During the preparation of this work, the authors used Microsoft 365 Copilot, based on the GPT-5 l, to assist programming, code-review, tests, validation-script, packaging, documentation, repository-organization, manuscript-structuring, language-refinement, and figure-planning. After using this tool, the authors reviewed and edited the content as needed and take full responsibility for the content of the published article

## ABBREVIATIONS

AUC: Area Under the Curve
DDA: Data Dependent Acquisition
DIA: Data Independent Acquisition
GRAVY: Grand Average of Hydrophathy
PEP: Posterior Error Probability
pI: Isoeletric point
ROC: Receiver-operating-characteristic

