## Supplementary Methods for "SeekNSee: A Rule-Based Framework for Prioritizing Proteotypic Peptides"

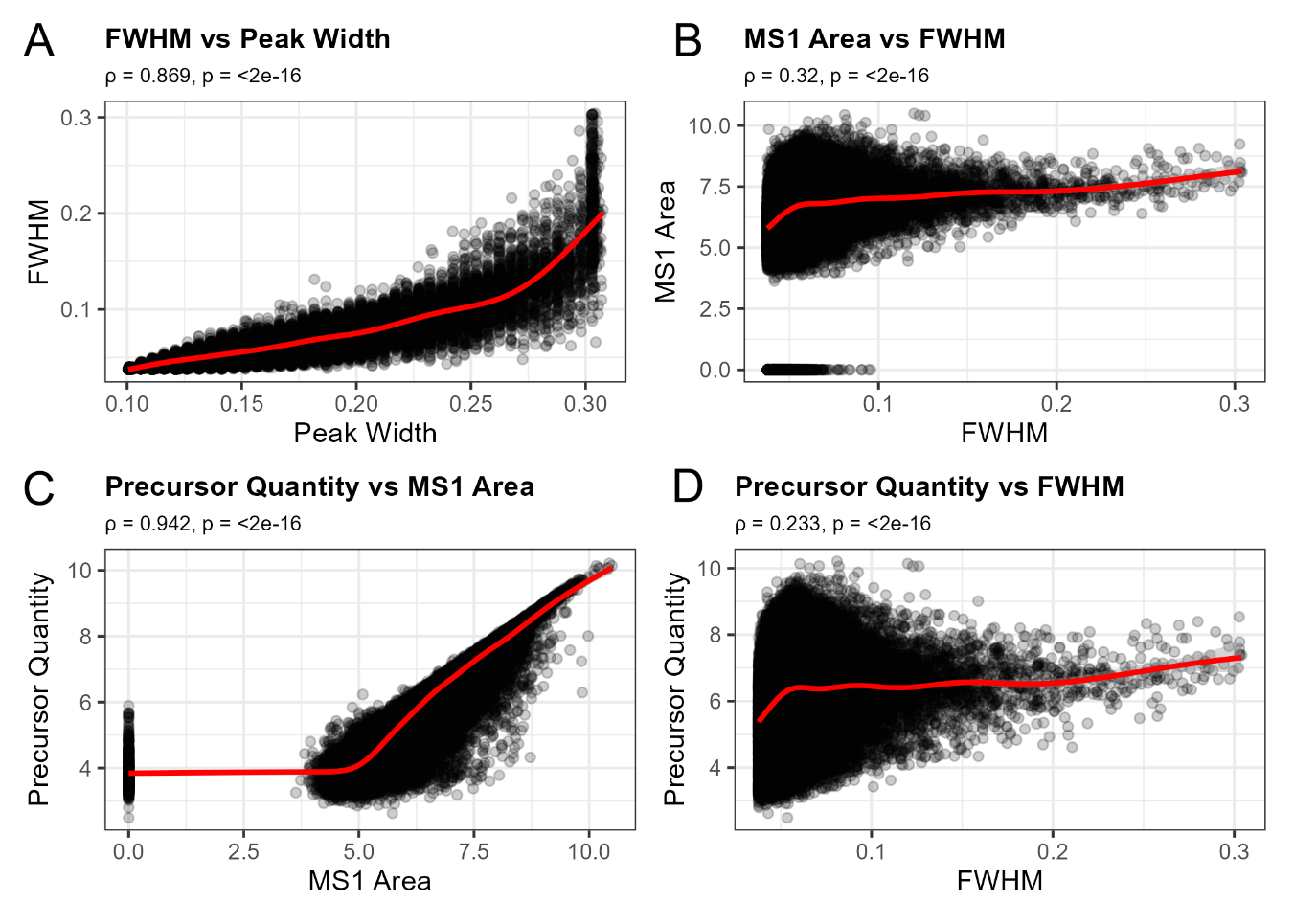

**Figure 1. Relationship between chromatographic peak dimensions and precursor quantification metrics.** Scatterplots depicting the relationships between **(A)** FWHM and peak width, **(B)** MS1 precursor area and FWHM, **(C)** Precursor quantity and MS1 precursor area, and **(D)** precursor quantity and FWHM. Red lines indicate LOESS fits with 95% confidence intervals. Spearman correlation coefficients (ρ) and corresponding *p*-values are shown in each panel.

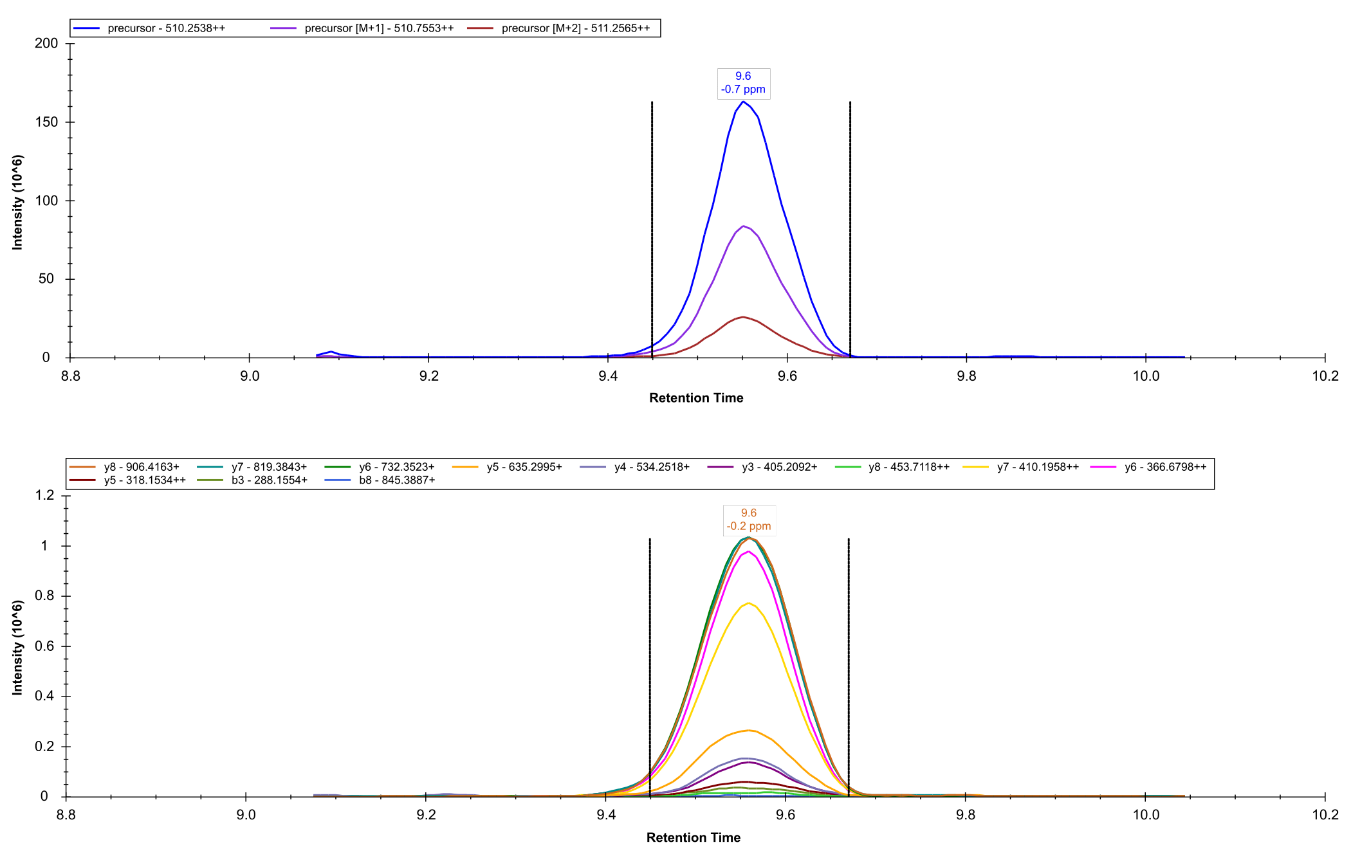

**Figure 2. Extracted ion chromatogram of ISSPTETER.** Precursor was selected as an illustration of high scoring precursor in terms of Empirical Quality, Quantity Quality, and Evidence reported by DIA-NN. Spectral libraries were imported into Skyline using DIA-NN compatibility for visualization.
